# *Prevotella copri* alleviates hyperuricemia via uricolysis and gut microbiota reconstruction

**DOI:** 10.64898/2026.05.04.722606

**Authors:** Yunpeng Yang, Meiling Yu, Linlin Yao, Peijun Yu, Yufei Huang, Xiaoman Yan

## Abstract

Despite the therapeutic promise of uric acid-metabolizing microbes for hyperuricemia, their application is hindered by the scarcity of highly efficacious strains. To identify uric acid-metabolizing microbes, we enriched gut microbiota from cynomolgus monkeys under uric acid-supplemented conditions. This selection increased the abundance of *Prevotellaceae_UCG-001* and *norank_f_Prevotellaceae*, implicating *Prevotella* species as key uric acid degraders. *In vitro* validation confirmed efficient uric acid consumption by two new *Prevotella* isolates and the reference strain *Prevotella copri* DSM 18205 (*P. copri*). Oral administration of *P. copri* in hyperuricemic mice reduced uric acid levels in serum and kidney tissues and alleviated renal fibrosis by suppressing the TGF-β/Smad pathway and downstream fibrogenic genes. Moreover, *P. copri* administration restored the depleted population of *Faecalibaculum*, and further assays demonstrated that *Faecalibaculum rodentium* directly metabolizes uric acid. Collectively, *P. copri* alleviates hyperuricemia via a dual mechanism: direct uric acid catabolism and enrichment of commensal uric acid-consuming bacteria. These findings establish *P. copri* as a promising live biotherapeutic agent for hyperuricemia and highlight microbial collaboration as a therapeutic strategy for metabolic diseases.

## Introduction

Hyperuricemia (HUA), characterized by elevated serum uric acid levels, poses significant health risks beyond its hallmark association with gout (Du *et al*, 2024) The most direct consequence is the crystallization of urate monosodium in joints and tissues, triggering intensely painful gouty arthritis(Yao *et al*, 2024). Chronic, uncontrolled hyperuricemia can lead to the formation of destructive tophi (urate crystal deposits) causing joint erosion and deformity(Jonsson *et al*, 2019). Crucially, hyperuricemia is a major risk factor for urolithiasis (kidney stones), particularly uric acid stones, causing renal colic and potential obstruction(Li *et al*, 2018). Furthermore, persistently high uric acid levels contribute to the development and progression of chronic kidney disease (CKD) through mechanisms like tubulointerstitial inflammation, fibrosis, and endothelial dysfunction(Li *et al*, 2021a; Wu *et al*, 2024). Evidence also links hyperuricemia to an increased risk of hypertension (Vareldzis *et al*, 2024; Wang *et al*, 2023), cardiovascular disease(Nishizawa *et al*, 2022; Waheed *et al*, 2021), and components of metabolic syndrome such as insulin resistance(Bahadoran *et al*, 2022; McCormick *et al*, 2021), and type 2 diabetes mellitus(Mortada, 2017).

Current hyperuricemia management prioritizes lowering serum uric acid (sUA) levels to prevent acute gout flares and mitigate long-term complications. Although foundational lifestyle modifications (e.g., dietary purine restriction, increased hydration, weight management) are essential, they frequently prove insufficient for achieving target sUA levels alone. Consequently, pharmacologic urate-lowering therapy (ULT) becomes necessary. However, the primary drug classes-xanthine oxidase inhibitors (e.g., allopurinol and febuxostat) (Hu & Brown, 2020; O’Dell *et al*, 2022), uricosuric agents(Li *et al*, 2023b), and uricase agents (e.g., pegloticase) (Seng Yue *et al*, 2008)-are associated with significant limitations and potential adverse effects that impact patient tolerance and long-term management. Therefore, developing new and safe strategies to lower serum uric acid levels is essential for the treatment of hyperuricemia.

The gut microbiome’s vast enzymatic repertoire, encoded by collective microbial genes, plays a crucial role in drugs and toxins metabolism, thus altering their bioavailability, activity, and toxicity(Dong *et al*, 2025; Manrique *et al*, 2024; Tian *et al*, 2023). Leveraging this potential, a promising strategy for treating hyperuricemia (HUA) focuses on screening the gut microbiota to identify bacterial strains harboring specific uric acid-metabolizing enzymes or efficient purine degradation pathways(Li *et al*, 2025; Liu *et al*, 2023; Liu *et al*, 2025; Tong *et al*, 2023). Until now, by isolating and characterizing these strains based on enzymatic activity or genetic markers (e.g., *ucd* genes), researchers had developed various new therapeutic strategies for hyperuricemia management(Wu *et al*, 2021; Zhao *et al*, 2022). However, the current scarcity of highly effective, mechanistically understood strains remains a major bottleneck for disease treatment.

In this study, we identify gut microbes with urate-metabolizing capacity through targeted enrichment. *Prevotella copri* emerges as a potent degrader, alleviating hyperuricemia and renal fibrosis in mice by directly consuming urate and suppressing the TGF-β/Smad signaling pathway and the expression of downstream fibrogenic genes. Furthermore, *P. copri* enriches for commensal *Faecalibaculum*, which also exhibits uricolytic activity, amplifying the therapeutic effect. These findings establish *P. copri* as a promising live biotherapeutic that targets hyperuricemia through a dual mechanism of direct and microbiota-enhanced urate clearance.

## Results

### Identification of uric acid-metabolizing gut microbes

Fecal samples from cynomolgus monkeys were homogenized in phosphate-buffered saline (PBS) supplemented with uric acid and subjected to anaerobic incubation for 15 hours. Following centrifugation, uric acid levels in the resultant supernatants were quantified (**Fig. 1A**). Comparative analysis showed that the fecal microbiota-supplemented group (Fm+uric acid) exhibited significantly higher uric acid metabolic activity than the PBS control (PBS+uric acid), which contained uric acid but lacked fecal microbes (**Fig. 1B**). To identify uric acid-metabolizing gut microbes, the fecal microbiome of Fm+uric acid and Fm+PBS groups were analyzed via 16S rRNA amplicon sequencing. Sequencing reads were rarefied to 24,601 reads per sample, yielding >99% average Good’s coverage (**fig. S1A**). Venn analysis revealed 529 amplicon sequence variants (ASVs) shared between the two groups, with 1,354 and 950 ASVs uniquely identified in the Fm+PBS and Fm+uric acid groups, respectively (**Fig. 1C**). The Fm+uric acid group showed significantly lower microbial richness (Ace and Chao indices) and diversity (Simpson index) relative to the Fm+PBS group (**Fig. 1D**). Principal coordinate analysis (PCoA) using Bray-Curtis distances further revealed distinct clustering of the Fm+uric acid microbial community from the control (Fm+PBS) (**Fig. 1E**). LEfSe (Linear Discriminant Analysis Effect Size) analysis identified distinct taxonomic enrichment patterns between groups: *norank_f_F082*, the phylum *Bacteroidota*, the order Pseudomonadales, *Treponema succinifaciens* DSM 2489, and *Acinetobacter* were enriched in the Fm+PBS group, whereas the genus *Treponema*, families *Spirochaetaceae* and *Rikenellaceae*, and *Prevotellaceae_UCG-001* were enriched in the Fm+uric acid group (**fig. S1B**).

**Fig. 1.**
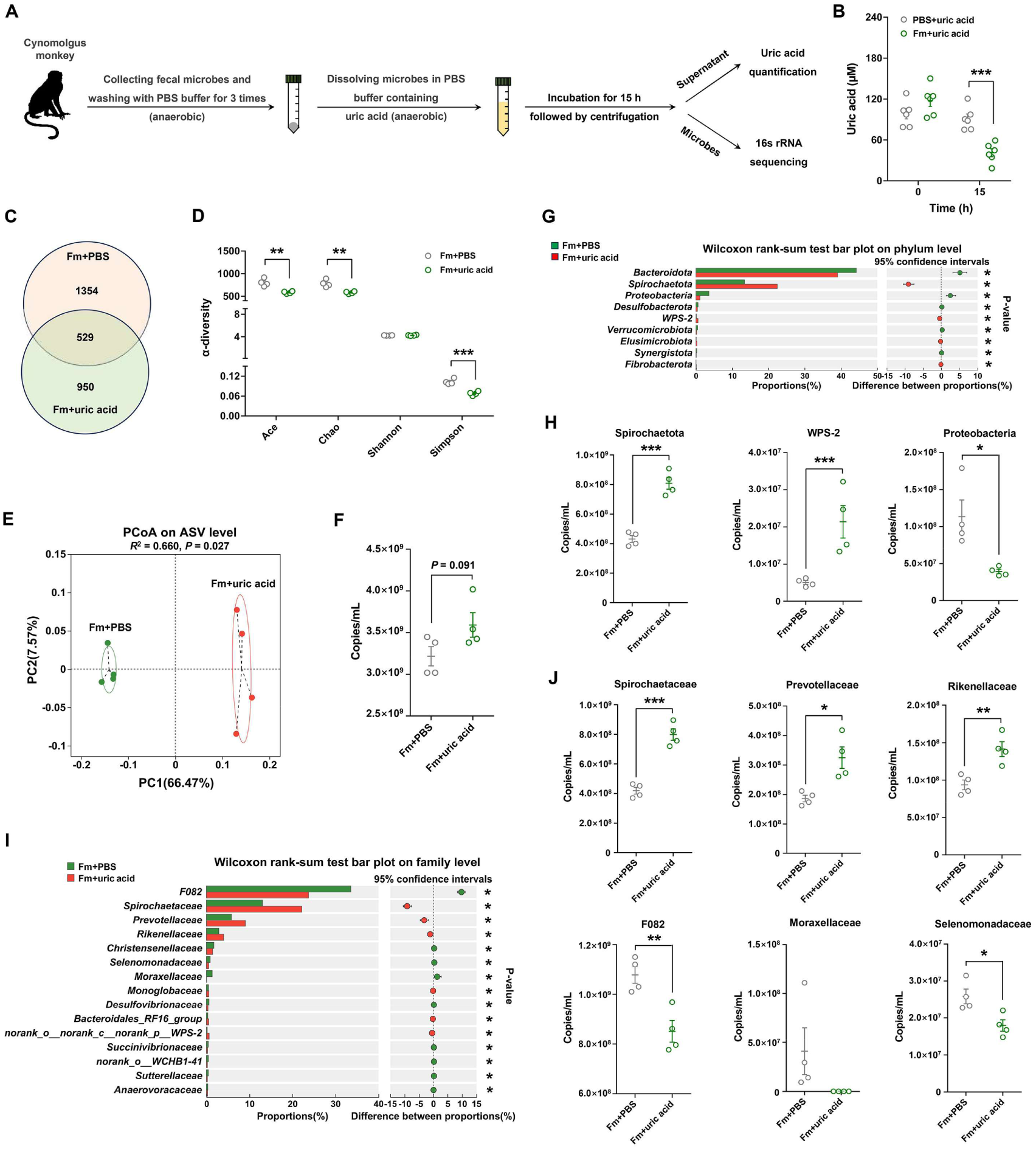
Uric acid supplementation alters the gut microbiome of cynomolgus monkey. (**A**) Schematic diagram depicting the strategy for isolating uric acid-metabolizing gut microbes. (**B**) Uric acid-metabolizing capacity of fecal microbiota. Two-tailed Welch’s t test, n = 6 for each group, presented as the mean ± SEM. (**C**) Venn diagram showing shared ASVs between the Fm+PBS and Fm+uric acid groups. (**D**) Gut microbiota richness (Ace and Chao indices) and diversity (Shannon and Simpson indices) in the two groups. (**E**) PCoA-based gut microbial differences of the two groups. (**F**) Total bacterial load quantification in the Fm+PBS and Fm+uric acid groups. (**G**) Relative abundance of gut microbiota at the phylum level. (**H**) Inferred absolute abundance of *Spirochaetota*, WPS-2, and *Proteobacteria* in both groups. (**I**) Relative abundance of gut microbiota at the family level. (**J**) Inferred absolute abundance of *Spirochaetaceae*, *Prevotellaceae*, *Rikenellaceae*, *F082*, and *Moraxellaceae* in both groups. Two-tailed Welch’s t test, n = 4 for each group, presented as the mean ± SEM. \**P* < 0.05, \*\**P* < 0.01, \*\*\**P* < 0.001.

To characterize gut microbial compositional shifts between the two groups, we profiled both relative and absolute abundances of the microbiota at the phylum, family, and genus levels. The Fm+uric acid group exhibited a significantly higher total bacterial load compared to the Fm+PBS control (**Fig. 1F**). Then, taxon-specific absolute abundances were calculated by multiplying the total fecal bacterial load by the relative abundance of each microbial taxon. At the phylum level, the Fm+uric acid group displayed significantly reduced abundance of *Proteobacteria*, whereas *Spirochaetota* and WPS-2 increased compared to the Fm+PBS control (**Fig. 1, G and H**). Family-level analysis showed enrichment of *Spirochaetaceae*, *Prevotellaceae,* and *Rikenellaceae* in the Fm+uric acid group, contrasting with depletion of norank_f_F082, *Moraxellaceae*, and *Selenomonadaceae* (**Fig. 1, I and J**). At the genus level, the Fm+uric acid group exhibited elevated abundances of *Treponema*, *Prevotellaceae_UCG-001*, *norank_f_Prevotellaceae*, and *unclassified_f_Prevotellaceae*, whereas *Acinetobacter* and *Pseudomonas* were reduced (**Fig. 2, A and B**).

**Figure 2.**
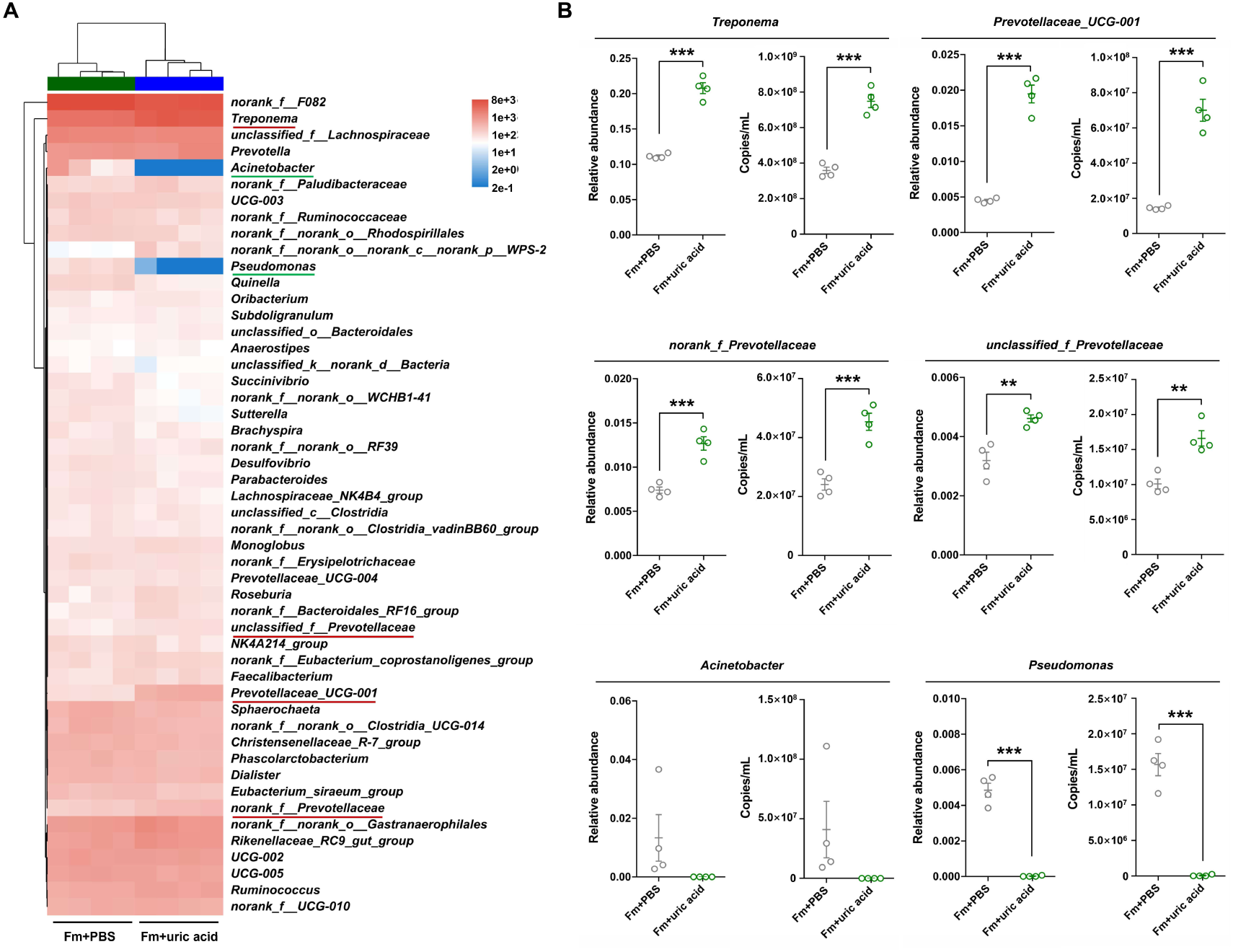
Uric acid supplementation alters the relative (A) and inferred absolute (B) abundance of gut microbiota at the genus level. Two-tailed Welch’s t test, n = 4 for each group, presented as the mean ± SEM.\**P* < 0.05, \*\**P* < 0.01, \*\*\**P* < 0.001.

The PICRUSt2-based heatmap analysis of MetaCyc pathways demonstrated distinct alterations in microbial functions between the Fm+uric acid and Fm+PBS groups. Fatty acid synthesis-related pathways (superpathway of fatty acid biosynthesis initiation (*E. coli*), (5Z)-dodec-5-enoate biosynthesis, palmitoleate biosynthesis I (from (5Z)-dodec-5-enoate), oleate biosynthesis IV (anaerobic), stearate biosynthesis II (bacteria and plants), and fatty acid elongation-saturated) were significantly downregulated in the Fm+uric acid group. Conversely, enhanced microbial functions in Fm+uric acid group were observed in three major categories: 1) Amino acid metabolism (L-lysine biosynthesis VI, superpathway of L-isoleucine biosynthesis I, superpathway of L-threonine biosynthesis, L-lysine biosynthesis III, superpathway of branched amino acid biosynthesis, L-isoleucine biosynthesis I (from threonine), L-valine biosynthesis, L-isoleucine biosynthesis IV, and L-isoleucine biosynthesis II); 2) Ribonucleotide biosynthesis (adenosine ribonucleotides de novo biosynthesis, 5-aminoimidazole ribonucleotide biosynthesis II, superpathway of 5-aminoimidazole ribonucleotide biosynthesis, and superpathway of pyrimidine ribonucleotides de novo biosynthesis); 3) Pyrimidine salvage pathway (superpathway of pyrimidine nucleobases salvage) (**fig. S2**).

Together, these findings demonstrate that uric acid supplementation drove significant restructuring of gut microbial composition, marked by a pronounced enrichment of Prevotellaceae-associated taxa.

### *Prevotella* microbes exhibits uric acid metabolizing capacity

The enrichment of *Prevotellaceae* under uric acid supplementation led us to hypothesize their capacity for uric acid metabolism. To test this, we isolated two *Prevotella* strains-*Prevotella* sp*. VZCB_s* iA622 and *Prevotella copri* CB7-from cynomolgus monkey feces (**Fig. 3A**). 16S rRNA sequence identity analysis confirmed both isolates shared high similarity (≥99%) with *Prevotella copri* DSM 18205 (**Fig. 3B**). We then cultured these three *Prevotella* strains in YCFAG medium supplemented with uric acid and collected the supernatants at 6 h to assess metabolic activity (**Fig. 3C**). Growth kinetics revealed that *Prevotella* sp*. VZCB_s* iA622 achieved significantly higher biomass yield than *P. copri* CB7 and *P. copri* DSM 18205, even in the absence of uric acid (**Fig. 3D**). Compared to uric acid-supplemented YCFAG medium without microbial inoculation (YCFAG group), the three *Prevotella* strains exhibited significant uric acid consumption (**Fig. 3E**). BLASTp-based comparative genomic analysis identified no reported uric acid-metabolizing enzyme orthologs in *Prevotella copri* DSM 18205(Li *et al*., 2025; Liu *et al*., 2023; Liu *et al*., 2025; Tong *et al*., 2023), suggesting a distinct uricolysis pathway in this strain (**fig. S3**).

**Fig. 3.**
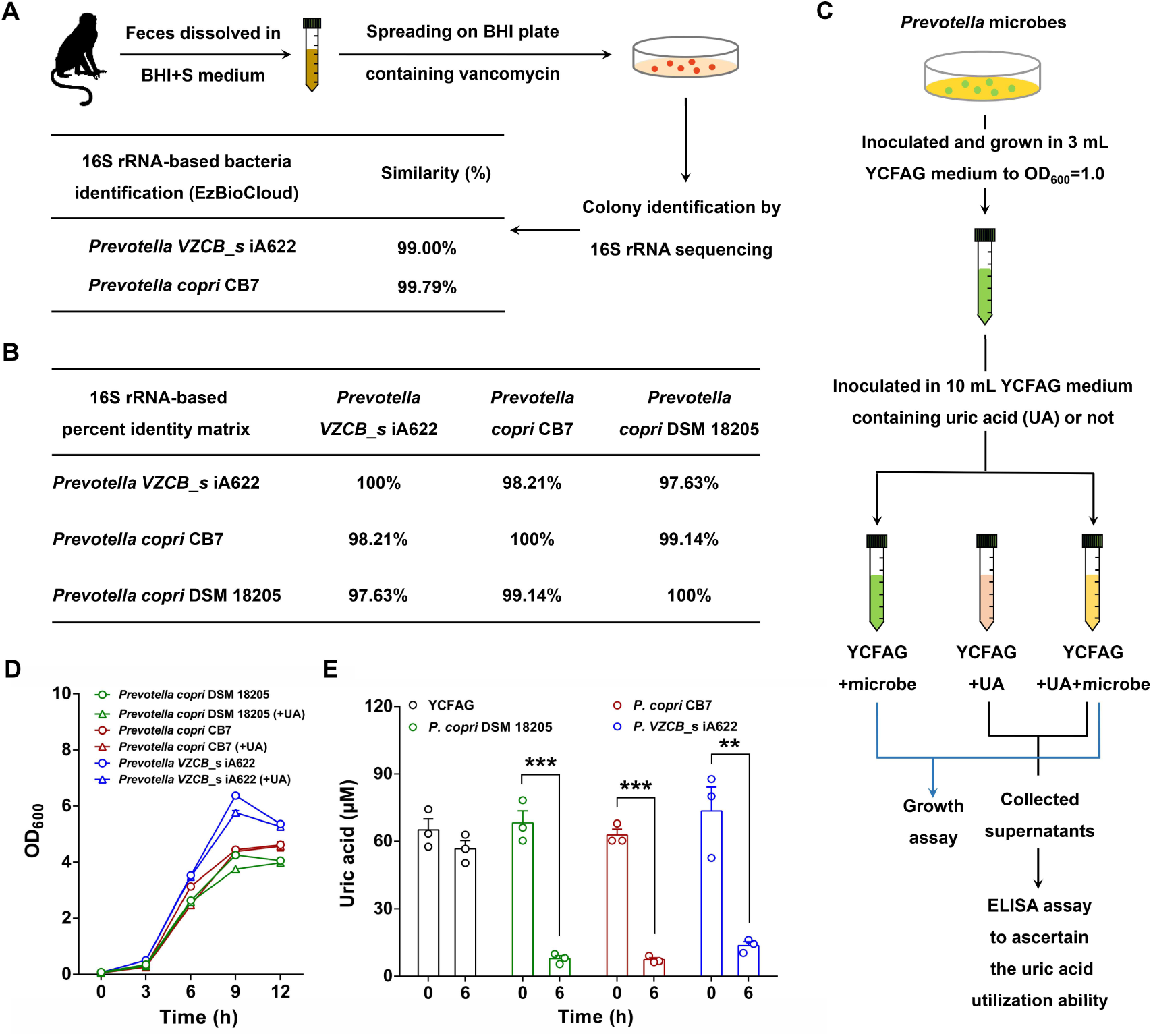
*Prevotella* microbes exhibit uric acid-metabolizing capacity. (**A**) Schematic diagram of the isolation strategy for *Prevotella* microbes from the fecal samples of cynomolgus monkey. (**B**) 16S rRNA-based percentage identity matrix among three Prevotella microbes. (**C**) Schematic diagram of the experimental design for assessing uric acid metabolism in three *Prevotella* microbes. (**D**) Growth kinetics of three Prevotella microbes in YCFAG medium supplemented with or without uric acid (UA). (**E**) The uric acid-metabolizing capacity of three *Prevotella* microbes at 6 h. Two-tailed Welch’s t test, n = 3 for each group, presented as the mean ± SEM.\**P* < 0.05, \*\**P* < 0.01, \*\*\**P* < 0.001.

### *Prevotella copri* ameliorates hyperuricemia and renal fibrosis through uric acid consumption and inhibition of the pro-fibrotic TGF-β/Smad pathway

To evaluate the *in vivo* uric acid-metabolizing capacity of *P. copri*, a hyperuricemia mouse model was induced by potassium oxonate and hypoxanthine co-administration to assess its therapeutic potential (**Fig. 4A**). Relative to the untreated control group (Control), hyperuricemic (HUA) mice exhibited significant body weight loss and reduced weight gain (**Fig. 4, B and C**). Notably, *Prevotella copri* intervention (HUA+*P. copri*) reversed hyperuricemia-associated growth retardation, suppressed serum uric acid accumulation, and reduced xanthine oxidase activity alongside two uremic toxin markers (blood urea nitrogen and creatinine) (**Fig. 4, B to G**). Moreover, renal amelioration was further demonstrated by decreased kidney-to-body weight ratio, restored tissue appearance, attenuated histopathology injury, and diminished intrarenal urate deposition in HUA+*P. copri* mice versus HUA controls (**Fig. 4, H to K**). Masson staining demonstrated significant collagen deposition in the kidney of HUA group mice compared to control group. This deposition was attenuated in mice receiving *P. copri* supplementation (HUA+*P. copri* group) (**Fig. 4, L and M**). qRT-PCR analysis revealed hyperactivation of the pro-fibrotic TGF-β/Smad pathway and increased expression of fibrogenic genes (*α-Sma* and *Timp1*) in the kidneys of HUA mice compared to Controls (**Fig. 4, N and O**). Oral administration of *P. copri* normalized both the TGF-β/Smad pathway activity and fibrogenic gene expression to Control levels (**Fig. 4, N and O**). Furthermore, *P. copri* treatment significantly reduced renal expression of uric acid transporters (*Npt1* and *Glut9*) and proinflammatory cytokines (*IL-1β*, *IL-6*, and *TNF-α*) relative to HUA controls (**fig. S4**).

**Fig. 4.**
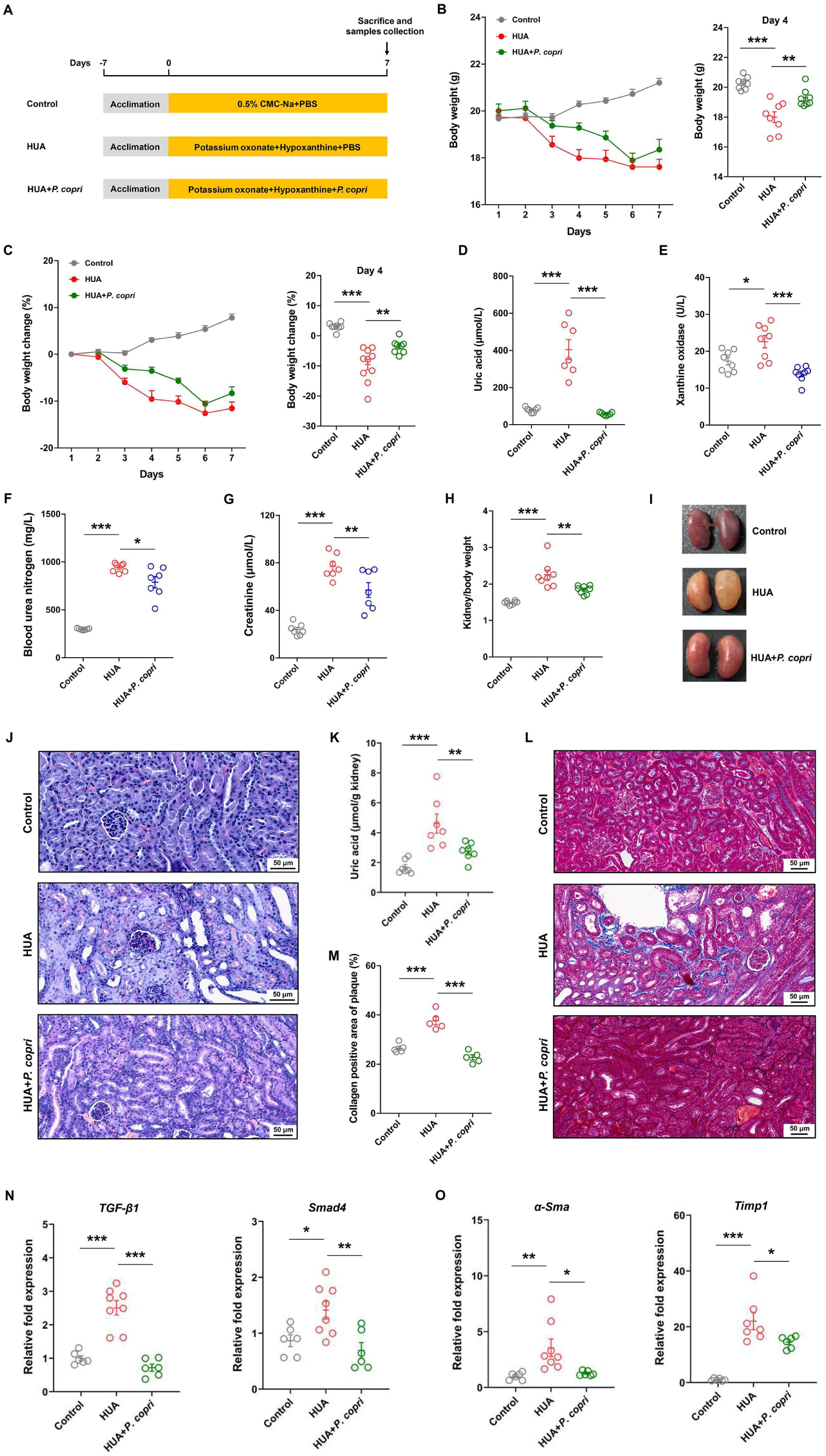
*P. copri* alleviates hyperuricemia and renal fibrosis via uricolytic activity and TGF-β/Smad pathway inhibition. (**A**) Schematic of hyperuricemia model induction and *P. copri* treatment protocol (n = 10 mice for each group). (**B**) Body weight of the three experimental groups. (**C**) Body weight change in the three groups. (**D**) Serum uric acid concentration in the three groups. (**E**) Serum xanthine oxidase activity in the three groups. (**F**) Blood urea nitrogen concentration in the three groups. (**G**) Serum creatinine concentration in the three groups. (**H**) Kidney-to-body weight ratio in the three groups. (**I**) Representative kidney appearance of three mice groups. (**J**) HE staining showing the renal histomorphology of three mice groups. (**K**) Kidney uric acid concentration in three groups. (**L**) Masson staining showing the renal collagen deposition in three groups. (**M**) Quantification of collagen-positive area in kidney sections of three mice groups. (**N**) Relative fold expression of *TGF-β1* and *Smad4* in the kidney of three mice groups. (**O**) Relative fold expression of *α-Sma* and *Timp1* in the kidney of three mice groups. One-way analysis of variance, n > 5 for each group, presented as the mean ± SEM.\**P* < 0.05, \*\**P* < 0.01, \*\*\**P* < 0.001.

Collectively, our data demonstrate that oral administration of *P. copri* alleviates hyperuricemia by reducing uric acid accumulation in serum and kidney tissues and attenuates kidney fibrosis by inhibiting the TGF-β/Smad signaling pathway, thereby suppressing the transcription of fibrogenic genes.

### *Prevotella copri* administration modulates gut microbiome composition of hyperuricemia mouse model

To assess the impact of *Prevotella copri* administration in the hyperuricemia mouse model, we performed 16S rRNA amplicon sequencing on duodenal, ileal, and colonic samples of three mice groups. Sequencing reads were rarefied to 25,362 per sample, achieving > 99% average Good’s coverage (**fig. S5A**). Venn diagram analysis identified 120, 54, and 302 core amplicon sequence variants (ASVs) shared across all mice groups in the duodenal, ileal, and colonic sites, respectively (**fig. S5B**). Although gut microbial diversity showed no significant differences among the three mouse groups at duodenal, ileal, and colonic sites, the gut microbial richness was reduced in the duodenum and ileum of both hyperuricemia groups (HUA and HUA+*P. copri*) compared to the Control group (**Fig. 5A**). Differently, no significant differences in gut microbial richness were observed in the colon of three mice groups (**Fig. 5A**). β-diversity analysis revealed significant compositional divergence between the Control group and two hyperuricemia groups (HUA and HUA+*P. copri*) across all intestinal sites (duodenum, ileum, colon) (**Fig. 5B**). Although the colonic microbiomes of the HUA and HUA+*P. copri* groups differed significantly, no significant differences were detected between these two groups in the duodenum or ileum (**Fig. 5B**).

**Fig. 5.**
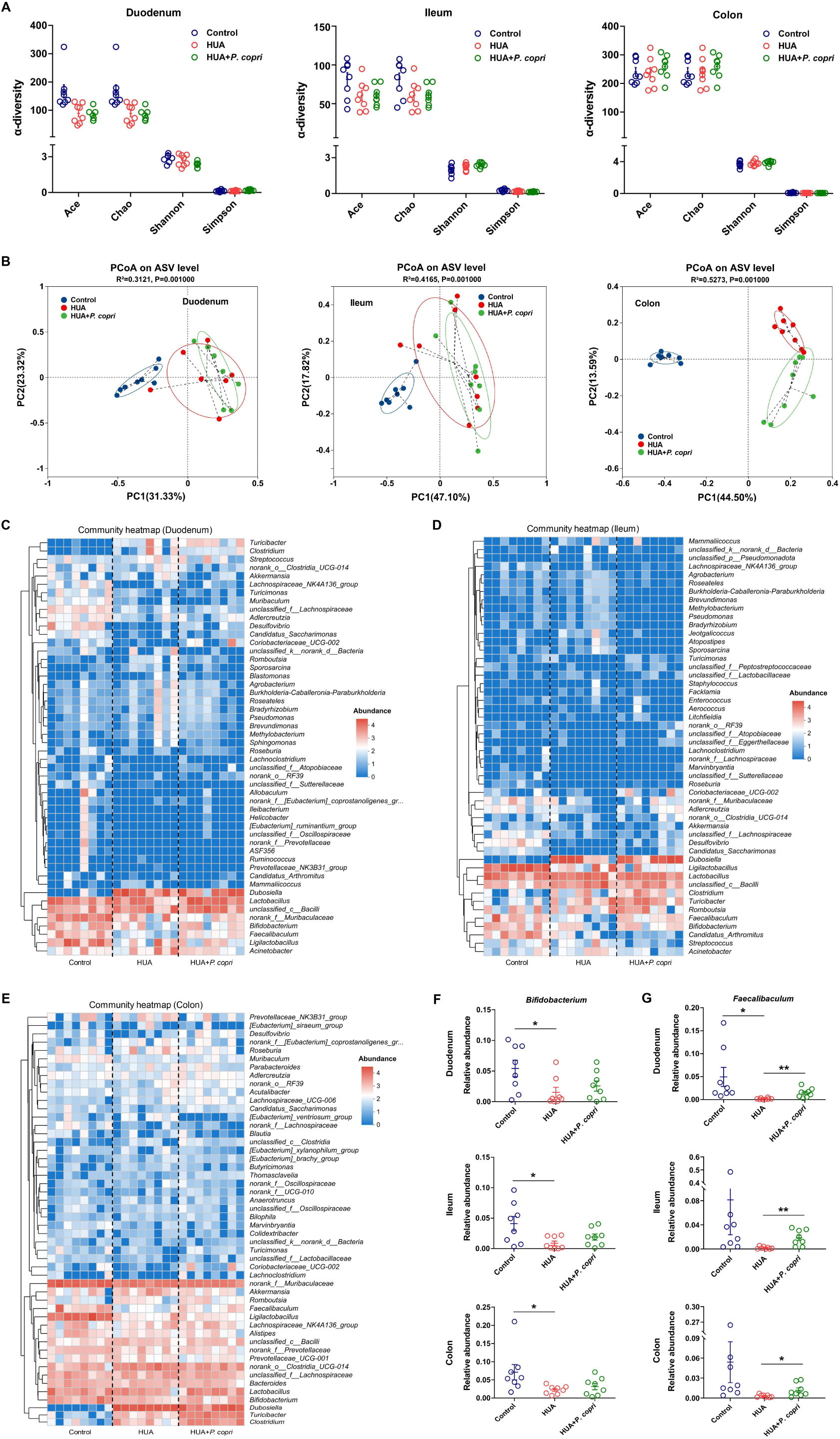
*Prevotella copri* administration modulates gut microbiota composition in the hyperuricemia mouse model. (**A**) Gut microbiota richness (Ace and Chao indices) and diversity (Shannon and Simpson indices) in the three mice groups at duodenum, ileum, and colon. (**B**) PCoA-based gut microbial differences between the three mice groups at duodenum, ileum, and colon. (**C-E**) Heatmap of top 50 genera in three mice groups at duodenum, ileum, and colon. (**F**) Relative abundance of *Bifidobacterium* between three mice groups at duodenum, ileum, and colon. (**G**) Relative abundance of *Faecalibaculum* between three mice groups at duodenum, ileum, and colon. One-way analysis of variance, n = 8 for each group, presented as the mean ± SEM.\**P* < 0.05, \*\**P* < 0.01, \*\*\**P* < 0.001.

LEfSe analysis identified microbiota signatures distinguishing the three mouse groups across intestinal niches. In the duodenum, *norank_f_Muribaculaceae*, *Ligilactobacillus*, *Faecalibaculum*, *Desulfovibrio*, and *norank_f_Eggerthellaceae* were enriched in Control group; *Dubosiella*, *Clostridium*, and *Turicibacter* dominated the HUA group; *Coriobacteriaceae_UCG-002* and *Lactobacillus intestinalis* were signature taxa in the HUA+*P. copri* group (**fig. S6A**). In the ileum, *Ligilactobacillus*, *Candidatus Arthromitus*, *Faecalibaculum*, *Desulfovibrio*, *norank_f_Muribaculaceae*, *Roseburia*, and *Adlercreutzia* were enriched in the Control group; *Streptococcus*, *Streptococcus danieliae*, *Sphingomonas*, *Bradyrhizobium*, *Jeotgalicoccus*, *Facklamia*, *Brevundimonas*, and *Agrobacterium* dominated the HUA group; The HUA+*P. copri* group was characterized by *Dubosiella*, *Turicibacter*, *Clostridium*, and *Coriobacteriaceae_UCG-002* (**fig. S6B**). In the colon, *Ligilactobacillus*, *norank_f_Muribaculaceae*, *Bifidobacterium*, *Faecalibaculum*, *norank_f_Prevotellaceae*, *Candidatus Arthromitus*, and *Streptococcus* were abundant in the Control group; *Dubosiella*, *norank_o_Clostridia_UCG-014*, and UCG-005 were enriched in the HUA group; *Turicibacter*, *Clostridium*, *Christensenellaceae_R-7_group* and *Coriobacteriaceae_UCG-002* dominated the HUA+*P. copri* group (**fig. S6C**). Based on these results, we generated heatmaps of the top 50 genera across three intestinal niches (duodenum, ileum, colon) to further examine the intergroup bacterial differences (**Fig. 5, C to E**). Compared to Controls, HUA mice exhibited decreased abundance of *Bifidobacterium* and *Faecalibaculum*, whereas supplementation with *P. copri* (HUA+*P. copri* group) increased the abundance of these two genera (**Fig. 5, F and G**).

Collectively, oral administration of *Prevotella copri* significantly restructured gut microbiota composition of three intestinal tract (duodenum, ileum, colon) in hyperuricemic mice.

### *Faecalibaculum rodentium* exhibits uricolytic activity

Given the increased abundance of *Bifidobacterium* and *Faecalibaculum* in hyperuricemic mice supplemented with *P. copri*, we hypothesized that these two genera might mediate uric acid metabolism. Spearman correlation analysis confirmed a significant negative correlation between *Faecalibaculum* abundance and serum uric acid levels across three intestinal niches (duodenum, ileum, colon), whereas *Bifidobacterium* exhibited no significant association (**Fig. 6A and fig. S7**), indicating that *Faecalibaculum* microbes might be the functional uric acid consumer. To test this hypothesis, we evaluated the uric acid-metabolizing capacity of *Faecalibaculum rodentium*. Specifically, the grown cells of *F. rodentium* were inoculated into PYG medium supplemented with uric acid (*F. rodentium*+uric acid) or not (*F. rodentium*) (**Fig. 6B**). The growth ability of *F. rodentium* was enhanced by uric acid supplementation (**Fig. 6C**). Consistent with this growth promotion, *F. rodentium* exhibited uric acid degradation activity when cultured in the presence of uric acid (*F. rodentium*+uric acid), compared to the bacteria-free control (PYG+uric acid) (**Fig. 6D**) Based on these findings, we conclude that *P. copri* administration could boost uric acid consumption by augmenting the populations of uric acid-consuming *Faecalibaculum* in the gut.

**Fig. 6.**
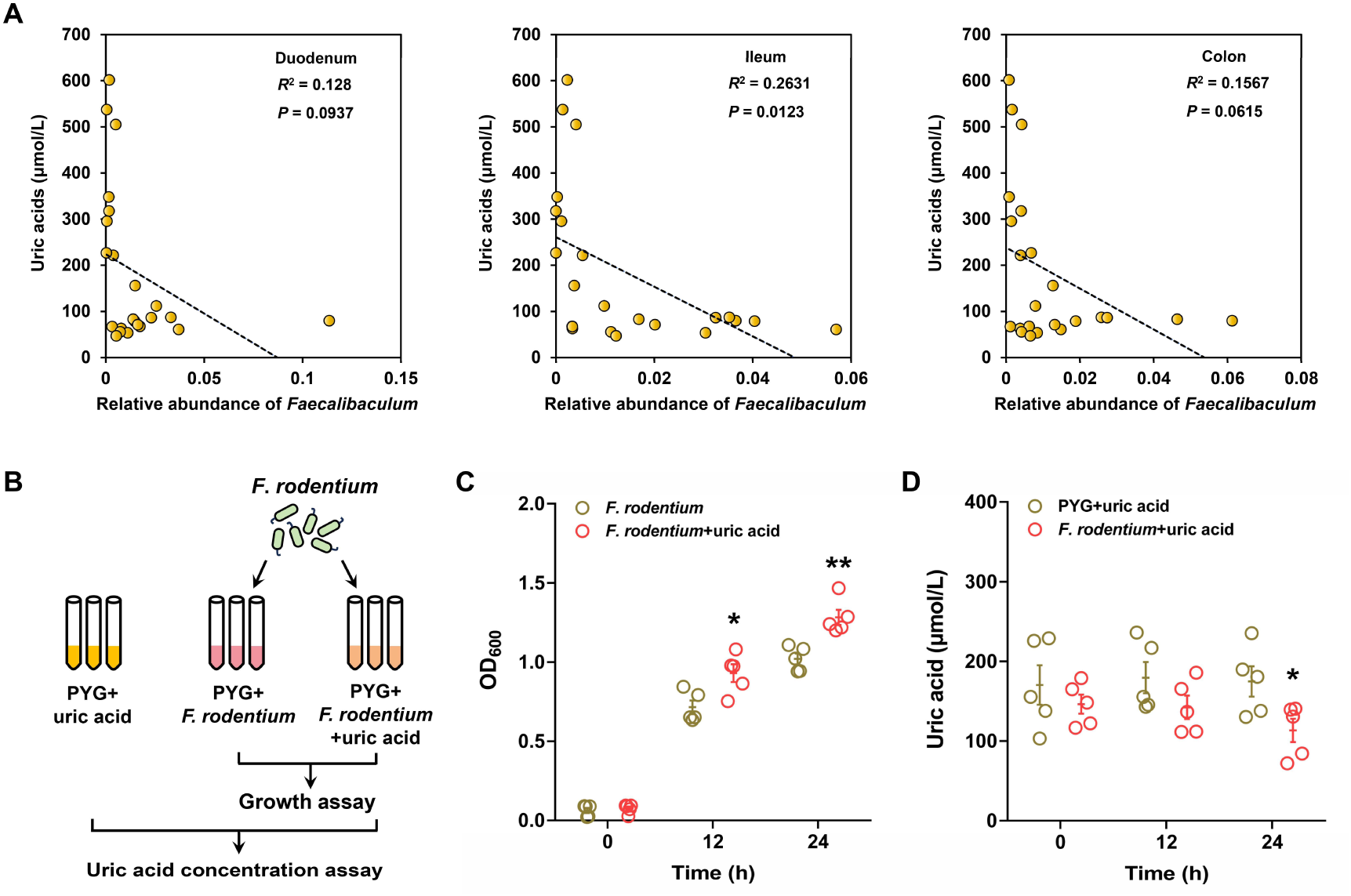
*Faecalibaculum rodentium* metabolizes uric acid directly. (**A**) Spearman-based correlation analysis between the relative abundance of *Faecalibaculum* and serum uric acid concentration. (**B**) Schematic diagram illustrating the strategy used for assaying the uric acid consumption ability. (**C**) The growth of *F. rodentium* in the PYG medium containing uric acid (*F. rodentium*+uric acid) or not (*F. rodentium*). (**D**) The uric acid metabolism ability of *F. rodentium*. Two-tailed Welch’s t test, n = 5 for each group, presented as the mean ± SEM.\**P* < 0.05, \*\**P* < 0.01, \*\*\**P* < 0.001.

## Discussion

Our study demonstrates that the gut microbiota represents a rich source of microbes capable of metabolizing uric acid, with potential therapeutic applications for hyperuricemia. By enriching microbial consortia from cynomolgus monkey feces through *in vitro* selection in uric acid-supplemented medium, we identified *Prevotella* as a highly efficient uric acid-metabolizing genus. Oral gavage of *P. copri* in a murine model of hyperuricemia significantly reduced systemic and renal uric acid levels and ameliorated kidney fibrosis, mechanistically linked to the inhibition of the TGF-β/Smad signaling pathway and downregulation of pro-fibrotic genes. Notably, *P. copri* administration-induced hyperuricemia amelioration was achieved via dual synergistic mechanisms: directly reducing systemic uric acid levels and ecologically restoring depleted populations of the key commensal uric acid consumer *Faecalibaculum* (**Fig. 7**). These findings highlight the potential of leveraging targeted microbial interventions to treat hyperuricemia and its associated renal complications.

**Fig. 7.**
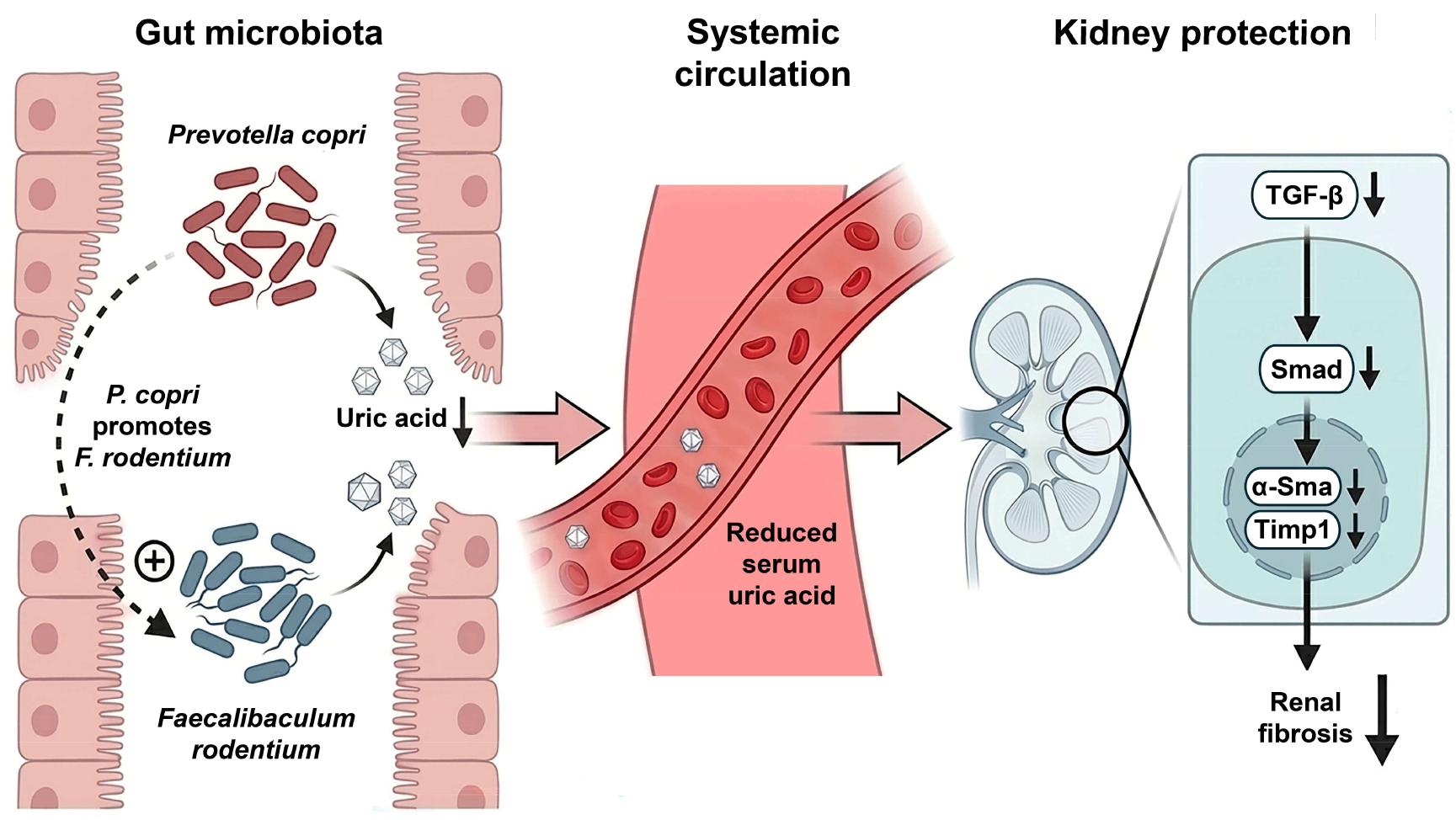
Schematic diagram illustrating *P. copri* administration-mediated hyperuricemia mitigation.

Beyond enriching *Prevotella*, uric acid supplementation significantly remodeled the cynomolgus monkey gut microbiome, increasing *Treponema* abundance, whereas reducing *Pseudomonas* (**Fig. 2B**). This suggests that the microbes of *Prevotella* and *Treponema* may function as uric acid consumers. However, given the pathogenic potential of most *Treponema* strains(Li *et al*, 2023a; Zhou *et al*, 2024), we focused exclusively on *Prevotella* microbes and functionally validated their uric acid-metabolizing capability (**Fig. 3E**). Furthermore, the divergent responses of *Prevotella* and *Pseudomonas* to uric acid exposure suggest that *Prevotella* may suppress *Pseudomonas* growth. This indicates a potential *Prevotella* administration-based therapeutic strategy for the treatment of diseases associated with *Pseudomonas* enrichment(Liao *et al*, 2022; Vidaillac & Chotirmall, 2021). The actual interaction between *Prevotella* and *Pseudomonas* microbes requires further validation through *in vitro* experiments.

Accumulating evidence reveals significant functional heterogeneity among *Prevotella copri* strains. For example, the *P. copri* strain isolated from swine feces activated chronic inflammatory responses in hosts via TLR4 and mTOR signaling pathways through its metabolites, thus increasing adiposity in pigs fed commercial formula diets(Chen *et al*, 2021); Three human fecal-derived *P. copri* strains (HF2123/clade A1, HF1478/clade A2, HF2130/clade C) could improve hyperglycemic mice metabolism by modulating gut microbial activity(Yang *et al*, 2024); Consortia containing the Bangladeshi *P. copri* strain PS131.S11 exerted anti-malnutrition effects by stimulating weight gain and reprogramming enterocyte energy metabolism(Chang *et al*, 2024). The functional divergence among *Prevotella copri* strains underlies their critical role in modulating host metabolic health.

In this work, two *Prevotella* strains capable of metabolizing uric acid were isolated from the fecal microbiome of cynomolgus monkey (**Fig. 3A**). Although these two isolates shared high percentage of 16S rRNA gene sequence and identical uric acid metabolic capability with the well-characterized *P. copri* strain DSM 18205, significant differences in growth kinetics were observed between *Prevotella VZCB_s* iA622 and the other two *Prevotella* strains (**Fig. 3, B, D, and E**). The conservation of uric acid catabolism across distinct *Prevotella* strains suggests that uric acid metabolism represents a conserved functional trait within this genus. Critically, the *P. copri* strain DSM 18205 used in this study exhibited a thoroughly evaluated safety profile(Verbrugghe *et al*, 2021), supporting its development as a next-generation probiotic for hyperuricemia prevention or management.

Despite the confirmed uric acid-metabolizing capability of *P. copri*, bioinformatic analysis targeting known uric acid metabolism-associated genes revealed the absence of characterized uricase or uric acid catabolism genes in its genome (**fig. S3**). This finding strongly suggests the existence of novel, yet unidentified, genetic determinants responsible for uric acid metabolism in this bacterium. To elucidate the uric acid metabolizing mechanism of *P. copri*, future work will employ transcriptomics, functional assays, and isotope tracing coupled with comprehensive metabolomics analyses to identify the key enzymes or pathways responsible for uric acid degradation and their downstream products, thereby clarifying the uric acid degrading mechanism in *P. copri*.

Oral administration of *P. copri* selectively altered the composition of the colonic microbiome, without affecting the duodenal or ileal microbiomes, in hyperuricemic mice (**Fig. 5B**). This site-specific effect is likely arises from the preferential colonization of *Prevotella* within the colon(Donaldson *et al*, 2016). Among the microbial shifts induced by *P. copri* administration, the abundance of *Faecalibaculum* significantly increased and was restored to baseline levels of the control group, which corroborated earlier findings(Chang *et al*., 2024). The underlying mechanism governing the interaction between *P. copri* and *Faecalibaculum* microbes merits further study. Although uric acid catabolism in *Faecalibaculum* had not been previously documented, our correlation analysis (linking *Faecalibaculum* abundance to serum uric acid concentration) and *in vitro* functional assays conclusively demonstrated the uric acid-metabolizing capability of *F. rodentium*. Future work will integrate bioinformatic, transcriptomics, and gene function analyses to elucidate the genetic basis and metabolic pathway of uric acid degradation in *F. rodentium*.

## Methods

### Reagents and tools table

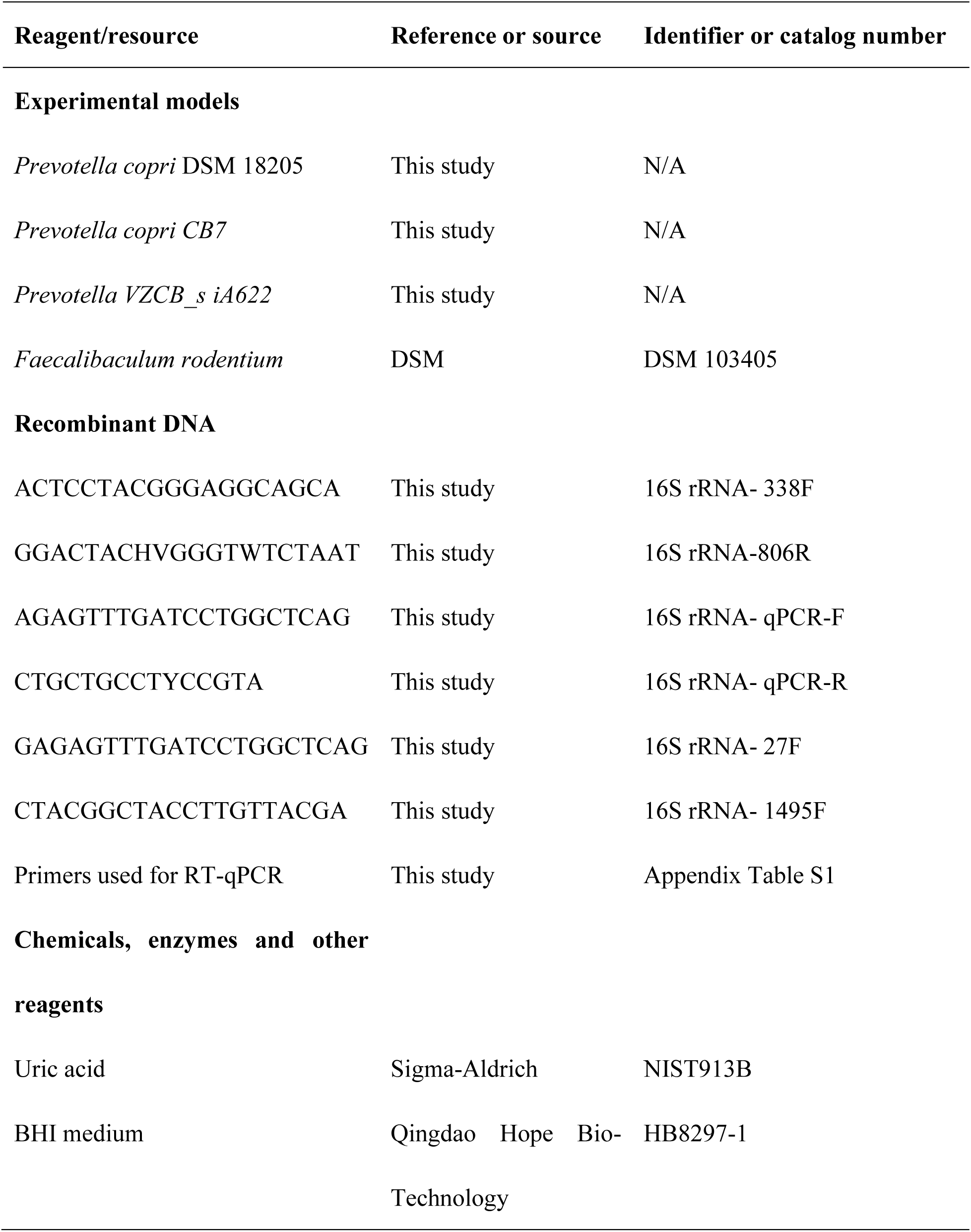
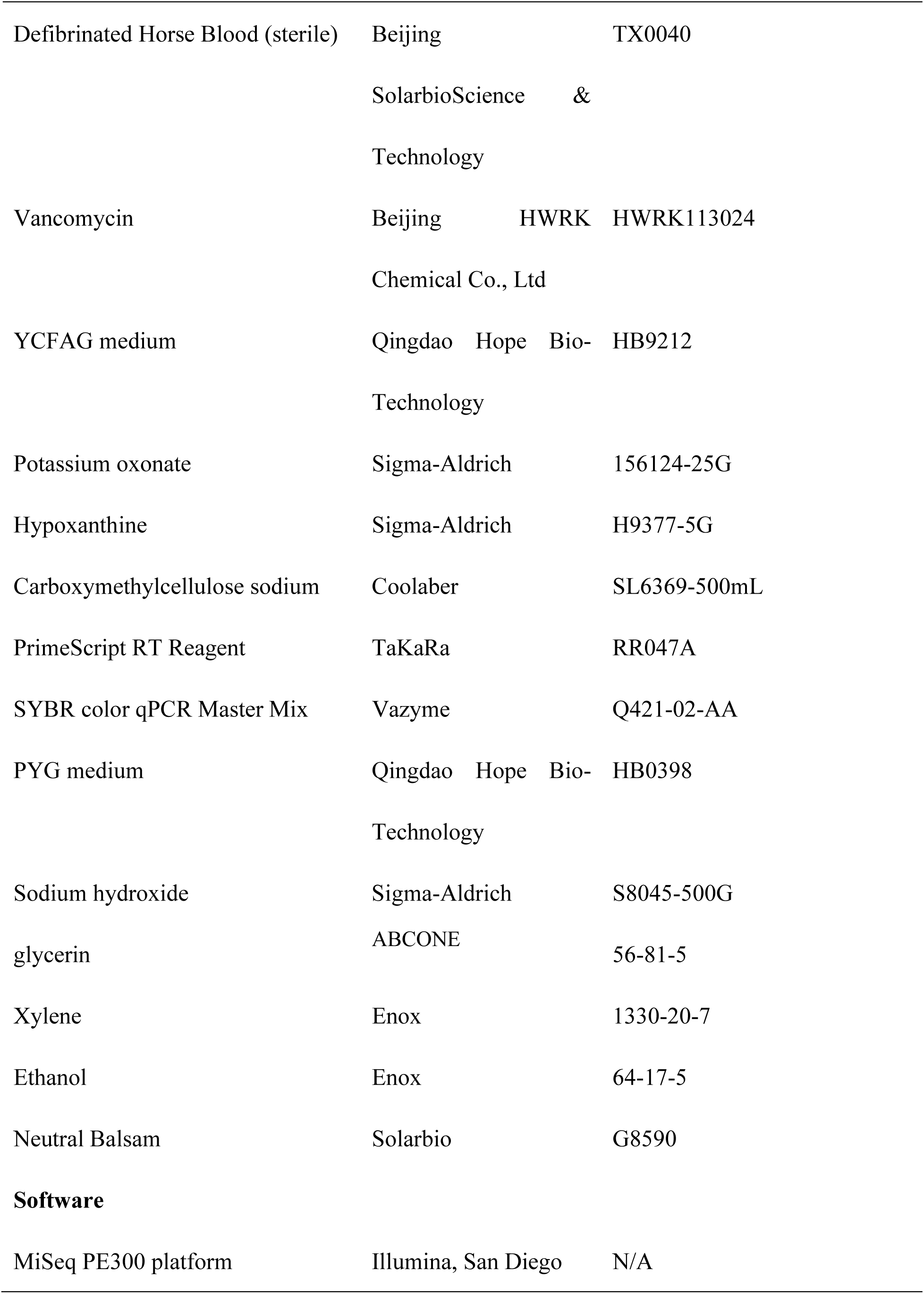
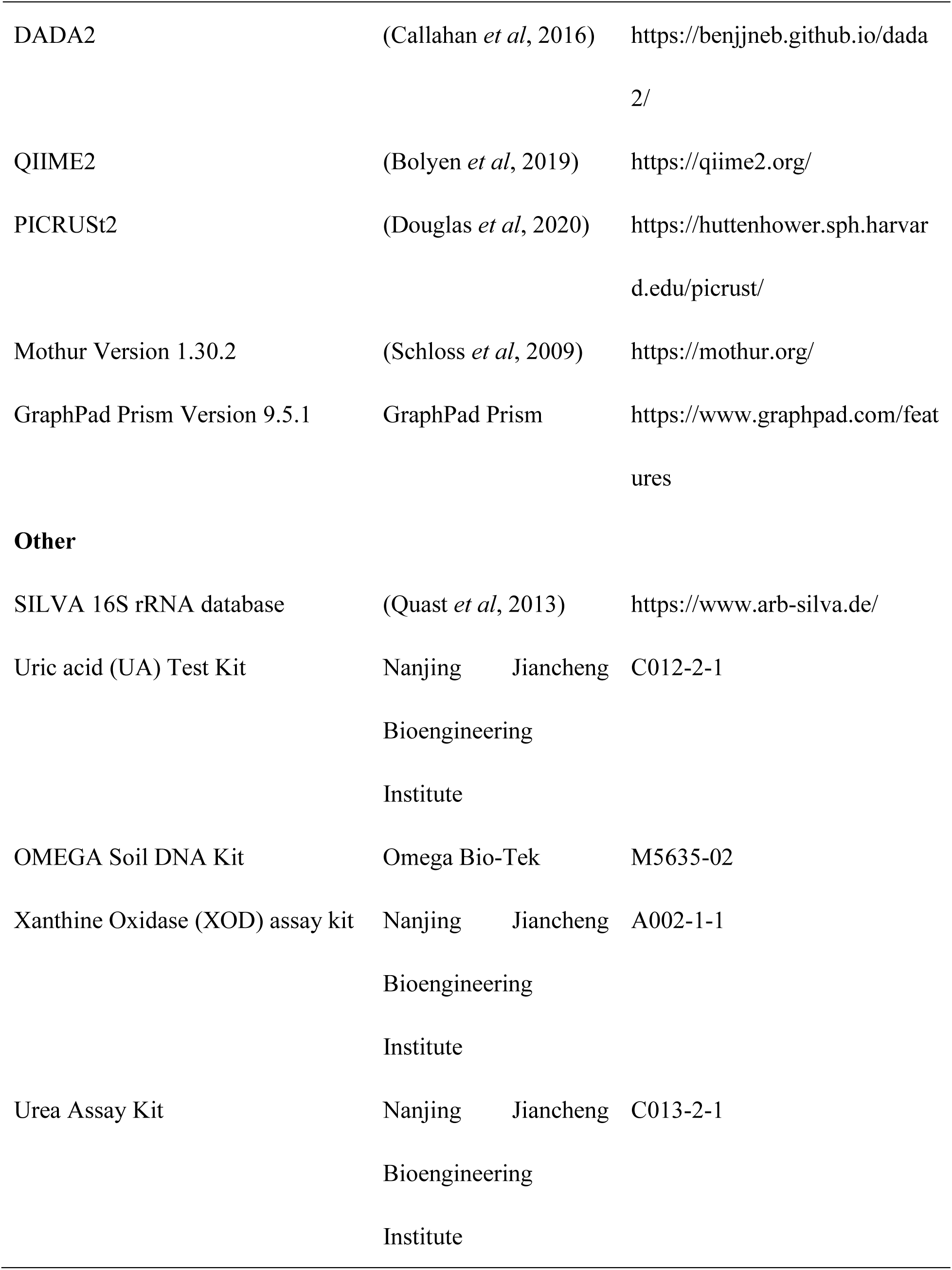
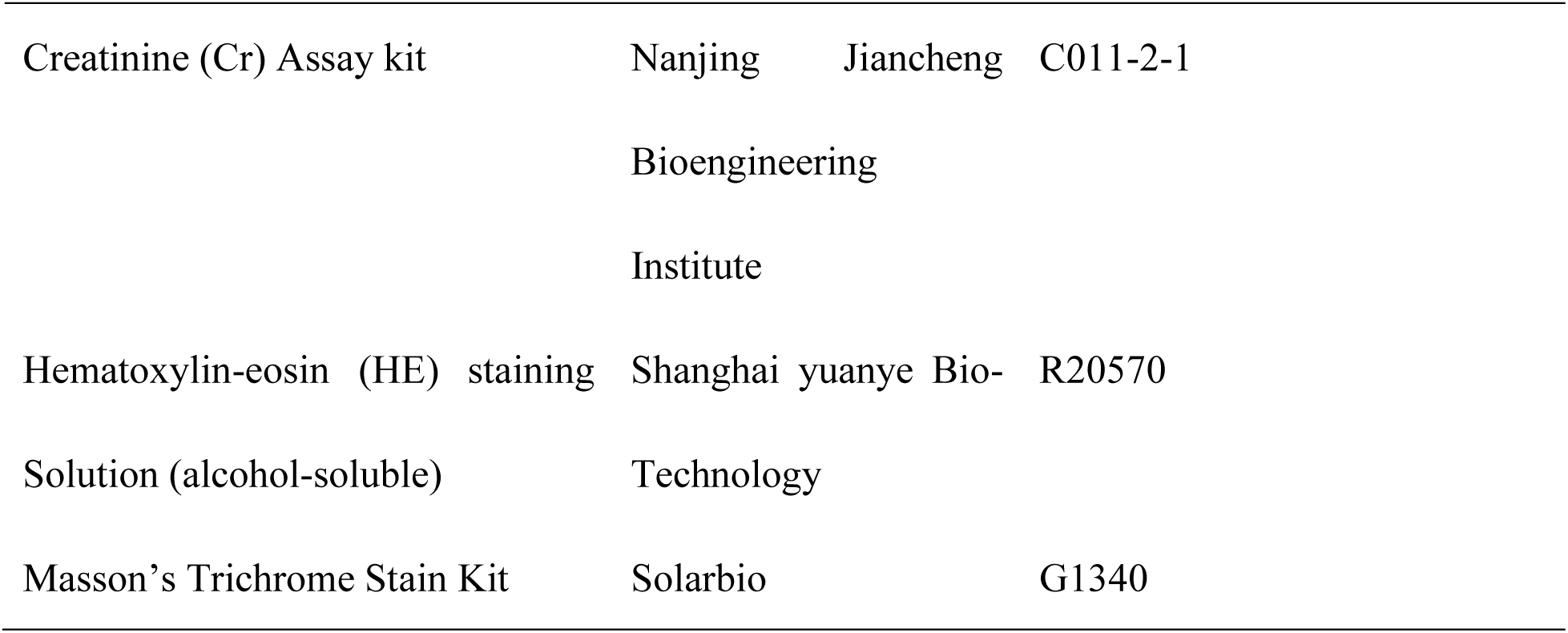

### Measurement of uric acid-metabolizing ability of Cynomolgus monkey-derived fecal microbes

The fecal samples of male Cynomolgus monkey were dissolved in PBS solutions (100 mg/mL), filtered by 70 μm cell strainer, and washed for 3 times. Then, the collected gut microbes were suspended in the PBS buffer containing 120 μM uric acid (Sigma-Aldrich, NIST913B) and incubated anaerobically for 15 h. After centrifugation, the concentration of uric acid in the supernatants were assayed by uric acid test kit (Nanjing Jiancheng Bioengineering Institute, C012-2-1).

### Ethics statement

The protocol for fecal sample collection is approved by the Animal Care Committee of Center for Excellence in Brain Science and Intelligence Technology, Chinese Academy of Sciences (No. NA-041-2022-R1).

### 16S rRNA gene amplicon sequencing

The genome extraction of gut microbes was performed by using the OMEGA Soil DNA Kit (Omega Bio-Tek, M5635-02). After quality examination, the genomic DNA concentration is assayed via NanoDrop ND-2000 spectrophotometer (Thermo Scientific Inc., USA). Then, the V3-V4 region of 16S rRNA gene is amplified by using the primers 338F (5’-ACTCCTACGGGAGGCAGCA-3’) and 806R (5’-GGACTACHVGGGTWTCTAAT-3’) as previously reported(Guo *et al*, 2025; Sun *et al*, 2024; Yang *et al*, 2025). After purification, the PCR products are paired-end sequenced with MiSeq PE300 platform (Illumina, San Diego, USA). The clean reads are de-noised using DADA2 (Callahan *et al*., 2016) to generate high-resolution amplicon sequence variants (ASVs), followed by taxonomic classification with the Naive Bayes consensus classifier implemented in QIIME2(Bolyen *et al*., 2019), referencing the SILVA 16S rRNA database (v138)(Quast *et al*., 2013). Then gut microbial function was predicted with PICRUSt2(Douglas *et al*., 2020).

All bioinformatic workflows were conducted on the Majorbio Cloud Platform. Alpha diversity metrics (e.g., Shannon, Chao1) were calculated using Mothur v1.30.2 (Schloss *et al*., 2009) to assess within-sample microbial richness and evenness. Beta diversity was evaluated via principal coordinate analysis (PCoA) derived from Bray-Curtis dissimilarity matrices, visualizing inter-sample community structural differences. The Venn diagram that generated with the R package VennDiagram was employed to illustrate shared and unique ASVs across different groups. To identify differentially abundant bacterial taxa, linear discriminant analysis effect size (LEfSe) (Segata *et al*, 2011)was applied, with a logarithmic LDA score threshold > 2.0 to prioritize biologically meaningful discriminators.

### 16S rRNA quantitative PCR

Total bacterial load in Fm+PBS and Fm+uric acid groups (15 h) was quantified via 16S rRNA quantitative PCR (qPCR) assay as previously reported(Yang *et al*, 2023). Amplification was performed using bacterial 16S rRNA-specific primers (forward: 5ʹ-AGAGTTTGATCCTGGCTCAG-3ʹ; reverse: 5ʹ-CTGCTGCCTYCCGTA-3ʹ) on a LightCycler 480 II system (Roche) under the following conditions: 95°C for 5 min (initial denaturation), followed by 45 cycles of 95°C for 10 s (denaturation), 60°C for 10 s (annealing), and 72°C for 10 s (extension). A standard curve constructed from a pMD18T vector harboring the full-length *E. coli* 16S rRNA gene was used for normalization in each qPCR run.

### Isolation of *Prevotella* microbes from the fecal sample of Cynomolgus monkey

The isolation of *Prevotella* microbes is performed as previously described with minor modifications(Li *et al*, 2021b). Briefly, fecal samples (100 mg) from male cynomolgus monkeys were homogenized in 1 mL of brain-heart infusion (BHI) medium (Qingdao Hope Bio-Technology, HB8297-1) supplemented with 10% fetal bovine serum (FBS) (Beijing SolarbioScience & Technology, TX0040). Serial dilutions of the homogenate were plated onto BHI-blood agar plates containing vancomycin (7 µg/mL) (Beijing HWRK Chemical Co., Ltd, HWRK113024). Plates were incubated under anaerobic conditions at 37°C for 48-72 h, after which isolated colonies were screened via single-colony PCR targeting the 16S rRNA gene with universal primers 27F (GAGAGTTTGATCCTGGCTCAG) and 1495R (CTACGGCTACCTTGTTACGA). Amplicons were Sanger-sequenced, and taxonomic assignment of *Prevotella* spp. was confirmed by BLASTn analysis.

### *In vitro* verification of uric acid-metabolizing ability of *Prevotella* microbes

Three *Prevotella* microbes (*Prevotella copri* DSM 18205, *Prevotella copri* CB7, and *Prevotella VZCB_s* iA622) were inoculated into 3 mL YCFAG medium (Qingdao Hope Bio-Technology, HB9212) and grown anaerobically at 37°C until OD_600_ reached 1.0. Then, the bacterial cells were inoculated into 10 mL YCFAG medium containing uric acid (60 μM) or not, and grown anaerobically at 37°C. The OD_600_ was assayed at 0, 3, 6, 9, and 12 h. The samples that collected at 0 and 6 h were centrifugated to collect the supernatant. The concentration of uric acid in the supernatants was assayed by uric acid test kit (Nanjing Jiancheng Bioengineering Institute, C012-2-1).

### Construction of hyperuricemia mice model

Acute hyperuricemia was induced in male C57BL/6J mice (4 weeks old) via daily intraperitoneal injection of potassium oxonate (PO, 300 mg/kg) (Sigma-Aldrich, 156124-25G)and hypoxanthine (HX, 300 mg/kg) (Sigma-Aldrich, H9377-5G)suspended in 0.5% (w/v) carboxymethylcellulose sodium (CMC-Na) (Coolaber, SL6369-500mL), following established methodologies(Xu *et al*, 2021). PO competitively inhibits uricase, blocking uric acid catabolism, whereas HX serves as a purine substrate to enhance uric acid synthesis via the xanthine oxidase pathway. Hyperuricemic induction was validated by quantifying serum uric acid levels using commercial assay kit (Nanjing Jiancheng Bioengineering Institute, C012-2-1), which revealed statistically significant increases compared to vehicle-treated controls receiving 0.5% CMC-Na alone.

### *P. copri* administration-based treatment of hyperuricemia

Hyperuricemic mice were divided into two experimental groups: 1) HUA group: Received daily oral administration of PBS containing 10% glycerol; 2) HUA+*P. copri* group: Received daily oral gavage of 1.0×10^8^ viable *P. copri* cells suspended in PBS with 10% glycerol (ABCONE, 56-81-5). Control mice received intraperitoneal injections of 0.5% CMC-Na followed by oral administration of PBS with 10% glycerol daily.

To evaluate the therapeutic efficacy of *P. copri* on hyperuricemia, mice from all three groups were sacrificed on day 7. Body weight was recorded daily to monitor weight changes. Serum and kidney uric acid levels were quantified using commercial assay kit (Nanjing Jiancheng Bioengineering Institute, C012-2-1). Similarly, xanthine oxidase activity, blood urea nitrogen (BUN), and serum creatinine concentrations were measured with commercial kits (Nanjing Jiancheng Bioengineering Institute, A002-1-1, C013-2-1, and C011-2-1). Kidney-to-body weight ratios were calculated at endpoint (day 7). Renal histopathology was assessed through hematoxylin and eosin (H&E) (Shanghai yuanye Bio-Technology, R20570) and Masson’s trichrome staining (Solarbio, G1340) following established protocols(Zhao *et al*, 2023).

### Quantitative real-time RT-PCR (qRT-PCR)

Renal tissue samples were collected from the Control, HUA, and HUA+*P. copri* groups. Total RNA was extracted using the TRIZOL method and reverse-transcribed into cDNA using the PrimeScript RT Reagent Kit (TaKaRa, RR047A). Quantitative real-time PCR (qRT-PCR) was performed with ChemQ SYBR color qPCR Master Mix (Vazyme, Q421-02-AA) under the following cycling conditions: initial denaturation at 95°C for 2 min; 40 cycles of 95°C for 15 s, 55°C for 20 s, and 72°C for 20 s. The relative expression levels of fibrogenic genes (*α-Sma* and *Timp1*), uric acid transporter genes (*Npt1* and *Glut9*), inflammatory genes (*IL-1β*, *IL-6*, and *TNF-α*) and TGF-β/Smad pathway genes in the three mice groups were normalized to that of GAPDH. All primers were provided in table S1.

### Assaying the uric acid consumption ability of *F. rodentium*

*F. rodentium* was cultured anaerobically in 3 mL of PYG medium (Qingdao Hope Bio-Technology, HB0398) at 37°C until OD_600_ reached 1.0. This culture was then inoculated (1% v/v) into fresh PYG medium (10 mL) supplemented with 200 μmol/L uric acid (Sigma-Aldrich, NIST913B) (*F. rodentium*+uric acid group). A control containing PYG medium with 200 μmol/L uric acid but no bacterial inoculum (PYG+uric acid group) was prepared in parallel. Both groups were incubated anaerobically at 37°C. After 12 h and 24 h of incubation, samples were centrifuged. The resulting supernatants from both the *F. rodentium*+uric acid and PYG+uric acid groups were collected, and uric acid concentration was quantified using a commercial uric acid assay kit (Nanjing Jiancheng Bioengineering Institute, C012-2-1).

### Evaluating the growth-promoting effect of uric acid on *F. rodentium*

*F. rodentium* was inoculated into 3 mL of PYG medium and grown anaerobically at 37°C until the OD_600_ reached 1.0. Subsequently, the culture was inoculated (1% v/v) into fresh 10 mL PYG medium either supplemented with 200 μmol/L uric acid (*F. rodentium*+uric acid group) or without uric acid (*F. rodentium* group). After anaerobic incubation at 37°C for 12 h and 24 h, the OD_600_ was measured to assess the effect of uric acid on *F. rodentium* growth.

### Quantification and statistical analysis

Data are presented as mean ± SEM. All statistical analyses were performed using GraphPad Prism (Version 9.5.1). For comparisons between two groups, statistical significance was assessed using an unpaired, two-tailed Student’s *t*-test. For comparisons involving more than two groups, one-way analysis of variance (ANOVA) was used, followed by Dunnett’s post hoc test for multiple comparisons.

## Data availability

The 16S rRNA gene amplicon sequencing data are available in the Sequence Read Archive of NCBI with Accession No. PRJNA1314121 and PRJNA1313829.

## Author contributions

Conceptualization, Yunpeng Yang; methodology, Meiling Yu, Linlin Yao, Peijun Yu, Yufei Huang, and Xiaoman Yan; investigation, Yunpeng Yang, Meiling Yu, Linlin Yao, and Peijun Yu; writing-original draft, Yunpeng Yang; writing-review and editing, Yunpeng Yang; funding acquisition, Yunpeng Yang and Yufei Huang; and supervision, Yunpeng Yang.

## Acknowledgements

This work was supported by the Basic Research Program of Jiangsu Province (BK20240910), the Natural Science Foundation of the Jiangsu Higher Education Institutions of China (24KJB230010), China Postdoctoral Science Foundation (2024M762747) and the Natural Science Foundation of Yangzhou City (YZ2024160). This work was also supported by the 111 Project D18007 and a Project Funded by the Priority Academic Program Development of Jiangsu Higher Education Institutions (PAPD).

## REFERENCES

Bahadoran Z, Mirmiran P, Kashfi K, Ghasemi A (2022) Hyperuricemia-induced endothelial insulin resistance: the nitric oxide connection. Pflügers Archiv - European Journal of Physiology 474: 83–98

Bolyen E, Rideout JR, Dillon MR, Bokulich NA, Abnet CC, Al-Ghalith GA, Alexander H, Alm EJ, Arumugam M, Asnicar F et al (2019) Reproducible, interactive, scalable and extensible microbiome data science using QIIME 2. Nature Biotechnology 37: 852–857

Callahan BJ, McMurdie PJ, Rosen MJ, Han AW, Johnson AJA, Holmes SP (2016) DADA2: High-resolution sample inference from Illumina amplicon data. Nature Methods 13: 581–583

Chang H-W, Lee EM, Wang Y, Zhou C, Pruss KM, Henrissat S, Chen RY, Kao C, Hibberd MC, Lynn HM et al (2024) Prevotella copri and microbiota members mediate the beneficial effects of a therapeutic food for malnutrition. Nature Microbiology 9: 922–937

Chen C, Fang S, Wei H, He M, Fu H, Xiong X, Zhou Y, Wu J, Gao J, Yang H et al (2021) Prevotella copri increases fat accumulation in pigs fed with formula diets. Microbiome 9: 175

Donaldson GP, Lee SM, Mazmanian SK (2016) Gut biogeography of the bacterial microbiota. Nature Reviews Microbiology 14: 20–32

Dong Z, Yu P, Li J, Zhou H, Li R, Wang S, Yang G, Nie Y, Liu L, Bian X et al (2025) Discovery of an ene-reductase initiating resveratrol catabolism in gut microbiota and its application in disease treatment. Cell Reports 44

Douglas GM, Maffei VJ, Zaneveld JR, Yurgel SN, Brown JR, Taylor CM, Huttenhower C, Langille MGI (2020) PICRUSt2 for prediction of metagenome functions. Nature Biotechnology 38: 685–688

Du L, Zong Y, Li H, Wang Q, Xie L, Yang B, Pang Y, Zhang C, Zhong Z, Gao J (2024) Hyperuricemia and its related diseases: mechanisms and advances in therapy. Signal Transduction and Targeted Therapy 9: 212

Guo L, Xu L, Nie Y, Liu L, Liu Z, Yang Y (2025) Murine gut microbial interactions exert antihyperglycemic effects. Isme j 19

Hu AM, Brown JN (2020) Comparative effect of allopurinol and febuxostat on long-term renal outcomes in patients with hyperuricemia and chronic kidney disease: a systematic review. Clinical Rheumatology 39: 3287–3294

Jonsson H, Aspelund T, Eiriksdottir G, Harris TB, Launer LJ, Gudnason V (2019) Hyperuricemia is associated with intermittent hand joint pain in a cross sectional study of elderly females: The AGES-Reykjavik Study. PLoS One 14: e0221474

Li CC, Chien TM, Wu WJ, Huang CN, Chou YH (2018) Uric acid stones increase the risk of chronic kidney disease. Urolithiasis 46: 543–547

Li H, Liu X, Lee MH, Li H (2021a) Vitamin C alleviates hyperuricemia nephropathy by reducing inflammation and fibrosis. J Food Sci 86: 3265–3276

Li J, Gálvez EJC, Amend L, Almási É, Iljazovic A, Lesker TR, Bielecka AA, Schorr EM, Strowig T (2021b) A versatile genetic toolbox for Prevotella copri enables studying polysaccharide utilization systems. The EMBO Journal 40: EMBJ2021108287

Li W, Yuan W, Huang S, Zou L, Zheng K, Xie D (2023a) Research progress on the mechanism of Treponema pallidum breaking through placental barrier. Microb Pathog 185: 106392

Li Y-J, Chen L-R, Yang Z-L, Wang P, Jiang F-F, Guo Y, Qian K, Yang M, Yin S-J, He G-H (2023b) Comparative efficacy and safety of uricosuric agents in the treatment of gout or hyperuricemia: a systematic review and network meta-analysis. Clinical Rheumatology 42: 215–224

Li Z, Meng W, Gao Z, Peng W, Hu Z, Zhang J, Wang Y, Wu X, Zhao Z, Zhang C et al (2025) A reductive uric acid degradation pathway in anaerobic bacteria. Life Metab 4: loaf031

Liao C, Huang X, Wang Q, Yao D, Lu W (2022) Virulence Factors of Pseudomonas Aeruginosa and Antivirulence Strategies to Combat Its Drug Resistance. Frontiers in Cellular and Infection Microbiology Volume 12–2022

Liu Y, Jarman JB, Low YS, Augustijn HE, Huang S, Chen H, DeFeo ME, Sekiba K, Hou BH, Meng X et al (2023) A widely distributed gene cluster compensates for uricase loss in hominids. Cell 186: 4472–4473

Liu Y, Zhou Z, Jarman JB, Chen H, Miranda-Velez M, Terkeltaub R, Dodd D (2025) Gut bacteria degrade purines via the 2,8-dioxopurine pathway. Nature Microbiology 10: 2291–2305

Manrique P, Montero I, Fernandez-Gosende M, Martinez N, Cantabrana CH, Rios-Covian D (2024) Past, present, and future of microbiome-based therapies. Microbiome Res Rep 3: 23

McCormick N, O’Connor MJ, Yokose C, Merriman TR, Mount DB, Leong A, Choi HK (2021) Assessing the Causal Relationships Between Insulin Resistance and Hyperuricemia and Gout Using Bidirectional Mendelian Randomization. Arthritis Rheumatol 73: 2096–2104

Mortada I (2017) Hyperuricemia, Type 2 Diabetes Mellitus, and Hypertension: an Emerging Association. Current Hypertension Reports 19: 69

Nishizawa H, Maeda N, Shimomura I (2022) Impact of hyperuricemia on chronic kidney disease and atherosclerotic cardiovascular disease. Hypertens Res 45: 635–640

O’Dell JR, Brophy MT, Pillinger MH, Neogi T, Palevsky PM, Wu H, Davis-Karim A, Newcomb JA, Ferguson R, Pittman D, et al (2022) Comparative Effectiveness of Allopurinol and Febuxostat in Gout Management. NEJM Evidence 1: EVIDoa2100028

Quast C, Pruesse E, Yilmaz P, Gerken J, Schweer T, Yarza P, Peplies J, Glöckner F (2013) The SILVA ribosomal RNA gene database project: improved data processing and web-based tools. Nucleic Acids Research 41: D590–D596

Schloss PD, Westcott SL, Ryabin T, Hall JR, Hartmann M, Hollister EB, Lesniewski RA, Oakley BB, Parks DH, Robinson CJ et al (2009) Introducing mothur: Open-Source, Platform-Independent, Community-Supported Software for Describing and Comparing Microbial Communities. Applied and Environmental Microbiology 75: 7537–7541

Segata N, Izard J, Waldron L, Gevers D, Miropolsky L, Garrett WS, Huttenhower C (2011) Metagenomic biomarker discovery and explanation. Genome Biology 12: R60

Seng Yue C, Huang W, Alton M, Maroli A, Waltrip R, Wright D, Marco M (2008) Population Pharmacokinetic and Pharmacodynamic Analysis of Pegloticase in Subjects With Hyperuricemia and Treatment-Failure Gout. Journal of clinical pharmacology 48: 708–718

Sun R, Yu P, Guo L, Huang Y, Nie Y, Yang Y (2024) Improving the growth and intestinal colonization of Escherichia coli Nissle 1917 by strengthening its oligopeptides importation ability. Metabolic engineering

Tian J, Li C, Dong Z, Yang Y, Xing J, Yu P, Xin Y, Xu F, Wang L, Mu Y et al (2023) Inactivation of the antidiabetic drug acarbose by human intestinal microbial-mediated degradation. Nat Metab 5: 896–909

Tong Y, Wei Y, Ju Y, Li P, Zhang Y, Li L, Gao L, Liu S, Liu D, Hu Y et al (2023) Anaerobic purinolytic enzymes enable dietary purine clearance by engineered gut bacteria. Cell Chem Biol 30: 1104–1114.e1107

Vareldzis R, Perez A, Reisin E (2024) Hyperuricemia: An Intriguing Connection to Metabolic Syndrome, Diabetes, Kidney Disease, and Hypertension. Curr Hypertens Rep 26: 237–245

Verbrugghe P, Brynjólfsson J, Jing X, Björck I, Hållenius F, Nilsson A (2021) Evaluation of hypoglycemic effect, safety and immunomodulation of Prevotella copri in mice. Scientific Reports 11: 21279

Vidaillac C, Chotirmall SH (2021) Pseudomonas aeruginosa in bronchiectasis: infection, inflammation, and therapies. Expert Review of Respiratory Medicine 15: 649–662

Waheed Y, Yang F, Sun D (2021) Role of asymptomatic hyperuricemia in the progression of chronic kidney disease and cardiovascular disease. Korean J Intern Med 36: 1281–1293

Wang C, Qin P, Liu Y, Wang L, Xu S, Chen H, Dai S, Zhao P, Hu F, Lou Y (2023) Association between hyperuricemia and hypertension and the mediatory role of obesity: a large cohort study in China. Rev Assoc Med Bras (1992) 69: e20220241

Wu F, Chen C, Lin G, Wu C, Xie J, Lin K, Dai X, Chen Z, Ye K, Yuan Y et al (2024) Caspase-11/GSDMD contributes to the progression of hyperuricemic nephropathy by promoting NETs formation. Cell Mol Life Sci 81: 114

Wu Y, Ye Z, Feng PY, Li R, Chen X, Tian XZ, Han R, Kakade A, Liu P, Li XK (2021) JL-3 isolated from “Jiangshui” ameliorates hyperuricemia by degrading uric acid. Gut Microbes 13

Xu L, Lin G, Yu Q, Li Q, Mai L, Cheng J, Xie J, Liu Y, Su Z, Li Y (2021) Anti-Hyperuricemic and Nephroprotective Effects of Dihydroberberine in Potassium Oxonate- and Hypoxanthine-Induced Hyperuricemic Mice. Frontiers in Pharmacology Volume 12–2021

Yang C, Lan R, Zhao L, Pu J, Hu D, Yang J, Zhou H, Han L, Ye L, Jin D et al (2024) Prevotella copri alleviates hyperglycemia and regulates gut microbiota and metabolic profiles in mice. mSystems 9

Yang Y, Yu P, Lu Y, Gao C, Sun Q (2023) Disturbed rhythmicity of intestinal hydrogen peroxide alters gut microbial oscillations in BMAL1-deficient monkeys. Cell Reports 42

Yang Y-P, Xu L-B, Lu Y, Wang J, Nie Y-H, Sun Q (2025) Dynamic alterations in bacterial and fungal microbiome and inflammatory cytokines following SRV-8 infection in cynomolgus monkeys. Zoological Research 46: 325–338

Yao T-K, Lee R-P, Wu W-T, Chen I-H, Yu T-C, Yeh K-T (2024) Advances in Gouty Arthritis Management: Integration of Established Therapies, Emerging Treatments, and Lifestyle Interventions. International Journal of Molecular Sciences 25: 10853

Zhao R, Li Z, Sun Y, Ge W, Wang M, Liu H, Xun L, Xia Y (2022) Engineered Escherichia coli Nissle 1917 with urate oxidase and an oxygen-recycling system for hyperuricemia treatment. Gut Microbes 14: 2070391

Zhao Y, Shao C, Zhou H, Yu L, Bao Y, Mao Q, Yang J, Wan H (2023) Salvianolic acid B inhibits atherosclerosis and TNF-α-induced inflammation by regulating NF-κB/NLRP3 signaling pathway. Phytomedicine 119: 155002

Zhou Y, Xie Y, Xu M (2024) Potential mechanisms of Treponema pallidum breaching the blood-brain barrier. Biomed Pharmacother 180: 117478

